# Discovery and structure–activity relationship analysis of 2-Pyridyl Thieno[3,2-d]pyrimidine derivatives as promising therapeutic candidates for the treatment of Buruli ulcer

**DOI:** 10.1101/2025.05.12.653443

**Authors:** Rodrigue Keumoe, Rufin Marie Toghueo Kouipou, Eugenie Aimée Madiesse Kemgne, Jean Baptiste Hzounda Fokou, Darline Dize, Patrick Valere Tsouh Fokou, Boniface Pone Kamdem, Adèle Honorine Enangue Makembe, Michelle Sidoine Nguembou Njionhou, Atanu Ghoshal, Abhijit Kundu, Benoît Laleu, Fabrice Fekam Boyom

## Abstract

*Mycobacterium ulcerans*, the bacterium causing Buruli ulcer (BU), can potentially develop resistance to existing antibiotics (rifampicin - clarithromycin/ moxifloxacin), underscoring the need for new antimycobacterial treatments. This study screened the Pathogen Box from Medicines for Malaria Venture (MMV) to identify *M. ulcerans* inhibitors. Four hit compounds were found, including the 2-(6-**methylpyridin-2-yl**)-N-(pyrimidin-4-yl)thieno[3,2-d]pyrimidin-4-amine **MMV688122** as a novel anti-*M. ulcerans* chemotype. Synthesis of structural analogues of **MMV688122** allowed the identification of 2-(4-**methylpyridin-2-yl**)-N-(pyrimidin-4-yl)thieno[3,2-d]pyrimidin-4-amine **MMV1578877** as the most potent, with submicromolar activity. Importantly, this analogue was non-cytotoxic up to 100 µM in human fibroblasts. Structure-activity relationship (SAR) studies indicated the crucial role of the methylpyridin-2-yl group in inhibiting *M. ulcerans* and the possibility to replace the thienopyrimidine core by a quinazoline. While MMV1578877 showed better metabolic stability than MMV688122, further improvement and testing in real-world *M. ulcerans* clinical isolates are still required. Further metabolite identification and SAR data should guide the optimization of this novel chemotype to enable *in vivo* testing.

**Author Summary:** Buruli ulcer is a neglected tropical disease that causes severe skin ulcers and long-term disability, mostly affecting people in remote African communities. Current treatments rely on antibiotics that are not always effective and may lead to resistance. In this study, we searched for new drug candidates by testing a library of compounds provided by the Medicines for Malaria Venture (MMV). We discovered a promising new chemical family that can kill the bacteria responsible for Buruli ulcer in the lab. One compound, in particular, showed strong activity without harming human cells. This compound also showed better stability and effectiveness. These findings bring us closer to developing a new, safer, and more effective treatment for Buruli ulcer.

## Introduction

Buruli ulcer caused by *Mycobacterium ulcerans* is a neglected tropical disease (NTD) reported in 33 countries in Africa, the Americas, Asia and the Western Pacific. This NTD is characterized by chronic, necrotizing lesions of the skin and soft tissue, leading to permanent disfigurement and disability if untreated. Moreover, this disease is featured by the development of painless swelling of the face, arms or legs (oedematous forms), that can be extended to bone infection, such as osteomyelitis [1,2]. The destruction of the skin is associated with the secretion of the macrolide toxin mycolactone by *M. ulcerans* that induces apoptosis and necrosis leading cell death. Mycolactone targets Sec61, a membrane protein complex that transports proteins to the endoplasmic reticulum, to induce immunosuppression and as a secondary effect, a stress response induction that can cause apoptosis and necrosis [3]. According to WHO, the first-line treatment for BU is a combination of rifampicin (10 mg/kg once daily) and clarithromycin (7.5 mg/kg twice daily). Moreover, the use of rifampicin (10 mg/kg once daily) along with moxifloxacin (400 mg once daily) is also recommended where clarithromycin is contraindicated. Notably, rifampicin has long been used as the main treatment for *Mycobacterium tuberculosis* infection and has gradually been extended to other *Mycobacterium* or non-tuberculous mycobacterial infections. This drug acts by disrupting the function of the DNA-dependent RNA polymerase [4]. Reported side effects associated with this medication include kidney and liver damage [5,6]. Other antibiotics in use, such as clarithromycin has been reported to induce pancreatitis, whereas moxifloxacin exhibits hepatic, cardiac, and gastrointestinal toxicities [7]. Thus, developing alternative and safer antimycobacterial agents is crucial to combating Buruli ulcer. Unfortunately, limited drug discovery efforts have been made against BU in the recent decade, resulting in the discovery of only a few countable hits and majorly repurposing anti-TB drug candidates such as TB47 and Q203 (Telacebec) [8,9]. Thienopyrimidines are heterocyclic compounds consisting of a fused thiophene and pyrimidine ring system, making them part of the broader class of fused heterocycles. Due to their structural versatility, thienopyrimidines have been explored in drug discovery, particularly for their antimicrobial, anticancer, and anti-inflammatory properties. In antibacterial drug development, thienopyrimidines have shown potential as inhibitors of key bacterial enzymes, including DNA gyrase, topoisomerases, and dihydrofolate reductase (DHFR), making them valuable scaffolds for designing new antibiotics [10]. This paper describes the potency and structure-activity relationship of a novel anti-*M. ulcerans* thienopyrimidine chemotype.

## Methods

### Chemical materials

The Pathogen Box was provided by MMV (Geneva, Switzerland). The analogues of MMV688122 were synthesized through MMV subcontract with TCG Lifesciences. Experimental procedures for the synthesis and characterization of MMV688122 and compounds 1-18 are detailed in the Supporting Information (S1&S2).

### Bacterial strain and culture conditions

*M. ulcerans* S4018, provided by the Biodefense and Emerging Infections Research Resources Repository (BEI Resources, Rockville, MD 20,852, USA), was cultured in Middlebrook 7H9 broth supplemented with 10% (v/v) oleic acid, albumin, dextrose and catalase (OADC) (0.06% oleic acid, 5% BSA, 2% dextrose, and 0.85% NaCl), glycerol (0.5% v/v) and tween 80 (0.05% v/v) and on Middlebrook 7H10 agar at 31°C. Media and chemicals were purchased from Sigma-Aldrich (Germany).

### Single point screening of compounds for antimycobacterial activity

To determine the ability of compounds to prevent the growth of *M. ulcerans in vitro*, the Pathogen Box compounds were screened in a 96-well plate format using the broth microdilution method [11]. Briefly, 98 µL of mycobacterial suspension at 10^6^ CFU/mL was treated with 2 µL of each compound at a single point concentration of 20 µM in a final volume of 100 µl in 96 well microtiter plates. Two hundred microliters of sterile distilled water were added to the surrounding wells to minimize evaporation during incubation. Sterility (distilled water), vehicle (0.4% DMSO) and positive (rifampicin) controls were included in each plate. The culture plates were sealed with plate seals (Sigma), wrapped in parafilm, and incubated at 31ºC for 8 days. After the incubation period, absorbance (ABS) was recorded at 630 nm using a microtiter plate reader (TECAN Infinite M200, Tecan, Männedorf, Switzerland). The recorded ABS values were used to calculate the percent of growth inhibition for each compound with regards to the growth control using equation below.

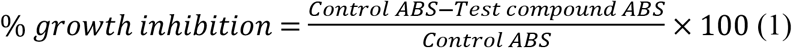

### Determination of the 50% inhibitory concentration (IC_50_)

Selected compounds showing ≥ 70% inhibition were tested for IC_50_ determination. Two-fold serial dilutions were performed to achieve test concentrations ranging from 20-0.039 µM for test compounds and 2-0.0078 µM for the positive control (rifampicin). Two microliters of each diluted sample were added to 98 µL of mycobacterium suspensions in a final volume of 100 µL per well. Sterility control containing medium alone and the vehicle control containing the solvent (0.4% DMSO) used for sample preparation and mycobacteria without test compounds were included in each plate. All experiments were performed in triplicate. Culture plates were incubated at 31°C for 8 days. Next, the ABS values were read at 630 nm using a microtiter plate reader (TECAN Infinite M200, Männedorf, Switzerland). The obtained ABS values were used to calculate the percentage of growth inhibition using MS Excel software as described above and the concentrations of test compounds that inhibited 50% of mycobacterial growth (IC_50_) relative to the vehicle control were determined using GraphPad Prism 5.0 software (Washington, DC, USA).

### Cytotoxicity assay

To determine the selectivity of the test compounds (MMV688122 and its analogues), their cytotoxicity was evaluated on human foreskin fibroblast HFF-1 cells (maintained in complete Dulbecco’s Modified Eagle’s Medium; DMEM) using the MTT [3-(4,5-dimethylthiazol-2-yl)-2,5-diphenyltetrazolium bromide] colorimetric assay. Briefly, 100 µL of human foreskin fibroblast HFF-1 cells (ATCC SCRC-1041) were seeded at 1×10^4^ cells/well in 96-well cell culture plates (CosStar, Washington, DC, United States of America) and incubated overnight to allow cell adhesion. Ten microliters of serially diluted compounds (100 - 1.5625 μM) were added and incubated for 48 h at 37°C under humidified atmosphere and 5% CO_2_. Ten microliters of dimethyl sulfoxide (DMSO; 0.4% *v/v*) was used as a negative control whereas podophyllotoxin (20 µM) served as a positive control. Twenty microliters of a stock solution of 2,5-diphenyl-2H-tetrazolium bromide (MTT) (5 mg/mL) were added to each well, gently mixed, and incubated for an additional 4 h. Next, after removing the culture medium, 100 µL of DMSO per well were added followed by incubation for 10 minutes at room temperature with gentle shaking. The ABS were then recorded at 570 nm using a microplate reader (TECAN-Infinite M200, Tecan, Männedorf, Switzerland). Compounds were tested in triplicate in two separate experiments. The median cytotoxic concentrations (CC_50_) of compounds were determined by analysis of concentration–response curves using the software GraphPad Prism 5.0 (Washington, DC, USA).

### LogD pH7.4 determination assay

The LogD pH7.4 assay was performed using a miniaturized shake flask method. A solution of a pre-saturated mixture of 1-octanol and phosphate buffered saline (PBS) (1:1, v/v) and the test compound (75 µM) was incubated at 25 °C with constant shaking (850 rpm) for 2 hours. The organic and aqueous phases were separated, and samples of each phase transferred to plate for dilution. The organic phase was diluted to 1000-fold and the aqueous phase was diluted 20-fold. The samples were quantitated using LC-MS/MS. The experiment was carried out in duplicate. Propranolol, amitriptyline and midazolam were used as reference standards.

### Aqueous solubility in phosphate buffered saline (PBS) pH7.4 assay

The kinetic solubility assay was performed using a miniaturized shake flask method. A solution of PBS and the test compound (200 µM) was incubated at 25 °C with constant shaking (600 rpm) for 2 hours. The samples were filtered using a multiscreen solubility filter plate. The filtrate was half diluted in acetonitrile. A five-point linearity curve was prepared in PBS:Acetonitrile (1:1, v/v) at 200, 150, 75, 25 and 2.5 µM. Blank, linearity and test samples (n = 2) were transferred to a UV readable plate and the plate was scanned for absorbance. Best fit calibration curves were constructed using the calibration standards and used to determine the test sample solubility. The experiment was carried out in duplicate. Diethylstilbestrol, haloperidol and sodium diclofenac were used as reference standards

### Metabolic stability study using human and mouse liver microsomes

A solution of the test compound in PBS (1 µM) was incubated in pooled human or mouse liver microsomes (0.5 mg/mL) for 0, 5, 20, 30, 45 and 60 minutes at 37 °C in the presence and absence of NADPH regeneration system (NRS). The reaction was terminated with the addition of ice-cold acetonitrile containing system suitability standard at designated time points. The sample was centrifuged (4200 rpm) for 20 minutes at 20 °C and the supernatant was half diluted in water and then analyzed by means of LC-MS/MS. The percentage of parent compound remaining, half-life (T1/2) and clearance (CLint,app) were calculated using standard methodology. The experiment was carried out in duplicate. Verapamil, diltiazem, phenacetin and imipramine were used as reference standards.

### Metabolic stability assay using cryopreserved mouse hepatocytes

A solution of the test compound in Krebs-Henseleit buffer solution (1 µM) was incubated in pooled mouse hepatocytes (1×106cells/mL) for 0, 15, 30, 45, 60, 75 and 90 minutes at 37 °C (5% CO2, 95% relative humidity). The reaction was terminated with the addition of ice-cold acetonitrile containing system suitability standard at designated time points. The sample was centrifuged (4200 rpm) for 20 minutes at 20 °C and the supernatant was half diluted in water and then analyzed by means of LC-MS/MS. The percentage of parent compound remaining, half-life (T1/2) and clearance (CLint,app) were calculated using standard methodology. The experiment was carried out in duplicate. Diltiazem, 7-ethoxy coumarin, propranolol and midazolam were used as reference standards.

### Metabolite identification studies

The *in vitro* metabolic stability of the most active antimycobacterial compound (MMV1578877) was evaluated in human and mouse liver microsomes (Xenotech, Lot #1410230 and 1510256, respectively) at 10 μM substrate and 1 mg/mL microsomal protein) The metabolic reaction was initiated by the addition of an NADPH-regenerating system and terminated at various time points over the incubation period by the addition of acetonitrile containing 150 ng/mL diazepam as an internal standard. Metabolite identification was performed using a Waters Xevo G2 QTOF coupled to a Waters Acquity UPLC using an Ascentis Express C8 column (50 × 2.1 mm, 2.7 µm) and detector, a positive electrospray ionisation under MSE mode 30 V. Mobile phase consisted of acetonitrile-water gradient with 0.05% formic acid at 4 min gradient cycle; 5 µL injection volume; 0.4 mL/min flow rate for metabolic stability and 6 min gradient cycle; 3 µL injection volume; 0.4 mL/min flow rate for metabolite identification. Metabolites were identified by extensive search using the Waters UNIFI software (Milford, Massachusetts, U.S.), accurate mass measurement and MS/MS scans.

## Results and Discussion

### Screening of the Pathogen Box for hit identification against *M. ulcerans*

The Pathogen Box was screened three times with *M. ulcerans* grown under identical culture conditions. A mean cut-off of 70% growth inhibition in the three independent screens of the compounds at 20 µM was applied as discrimination criterion of compounds with antimycobacterial activity. As a result, 37 hit compounds were identified and subsequently evaluated in concentration-response assays.

### Hit confirmation

The selected 37 hits were re-assayed using concentration-response tests to determine their median inhibitory concentrations (IC_50_). Four Pathogen Box compounds showed promising activity with IC_50_ values ranging from 0.59 µM to 3.34 µM. Compound MMV688775 (rifampicin, IC_50_ 0.59 µM) was the most active followed by MMV688122 (2-(6-methylpyridin-2-yl)-N-(pyrimidin-4-yl)thieno[3,2-d]pyrimidin-4-amine, IC_50_ 1.59 µM) and MMV688327 (radezolid, IC_50_ 1.6 µM), and lastly MMV688756 (sutezolid, IC_50_ 3.34 µM) (Table 1). The activity of rifampicin (MMV688775), which is a known antimycobacterial drug, has the merit of validating our tests and gives a reference value to compare the activities.

**Table 1.**
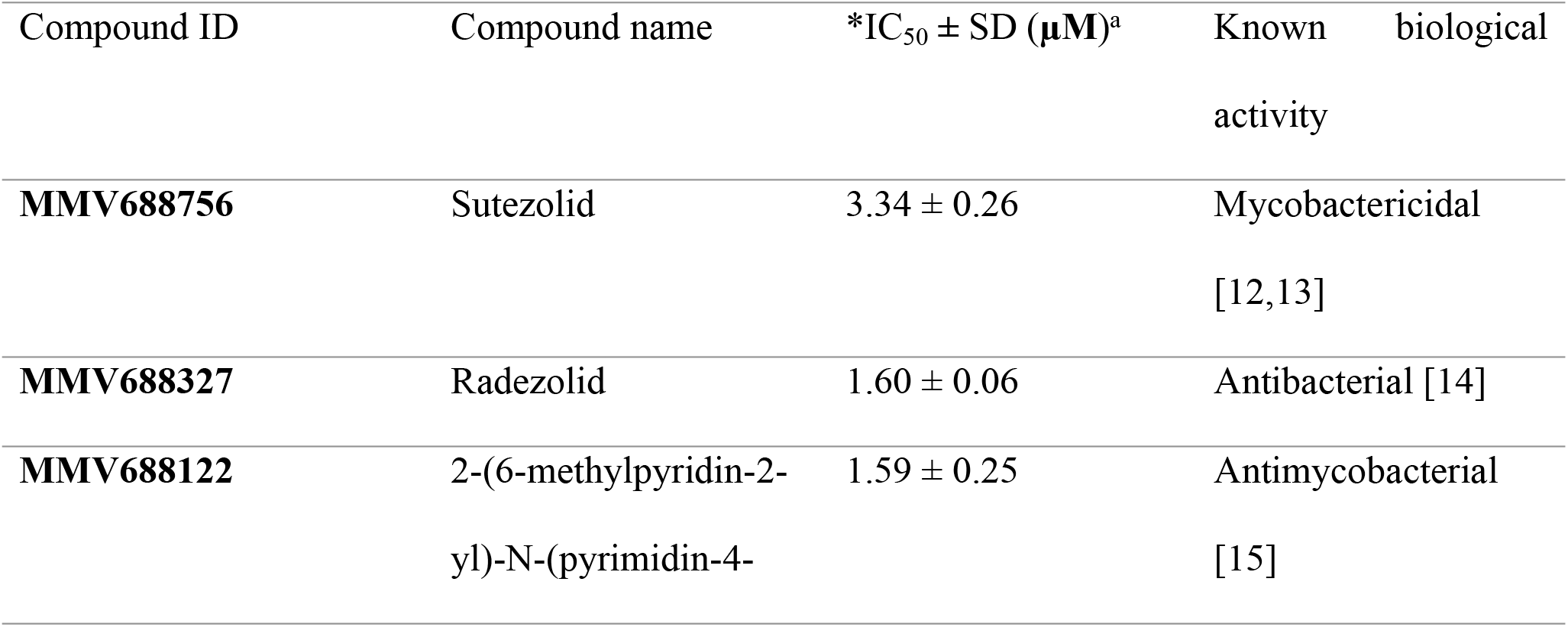
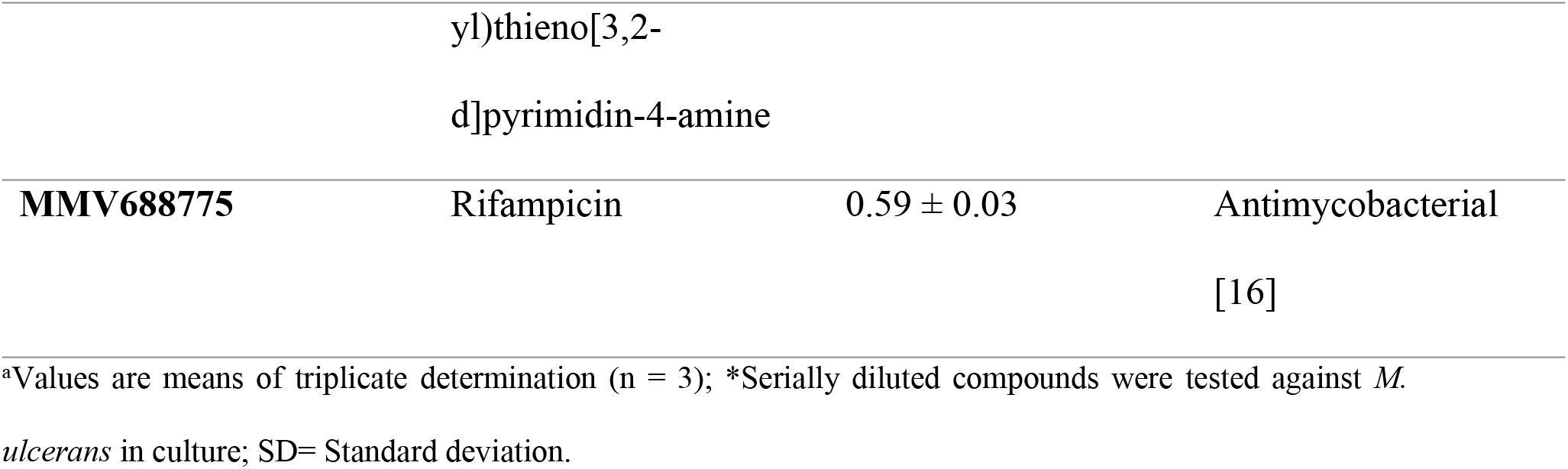
Inhibitory potential (IC_50_s) of anti-*M. ulcerans* compounds from the MMV Pathogen Box and their previously reported biological activities.

**Table 2.**
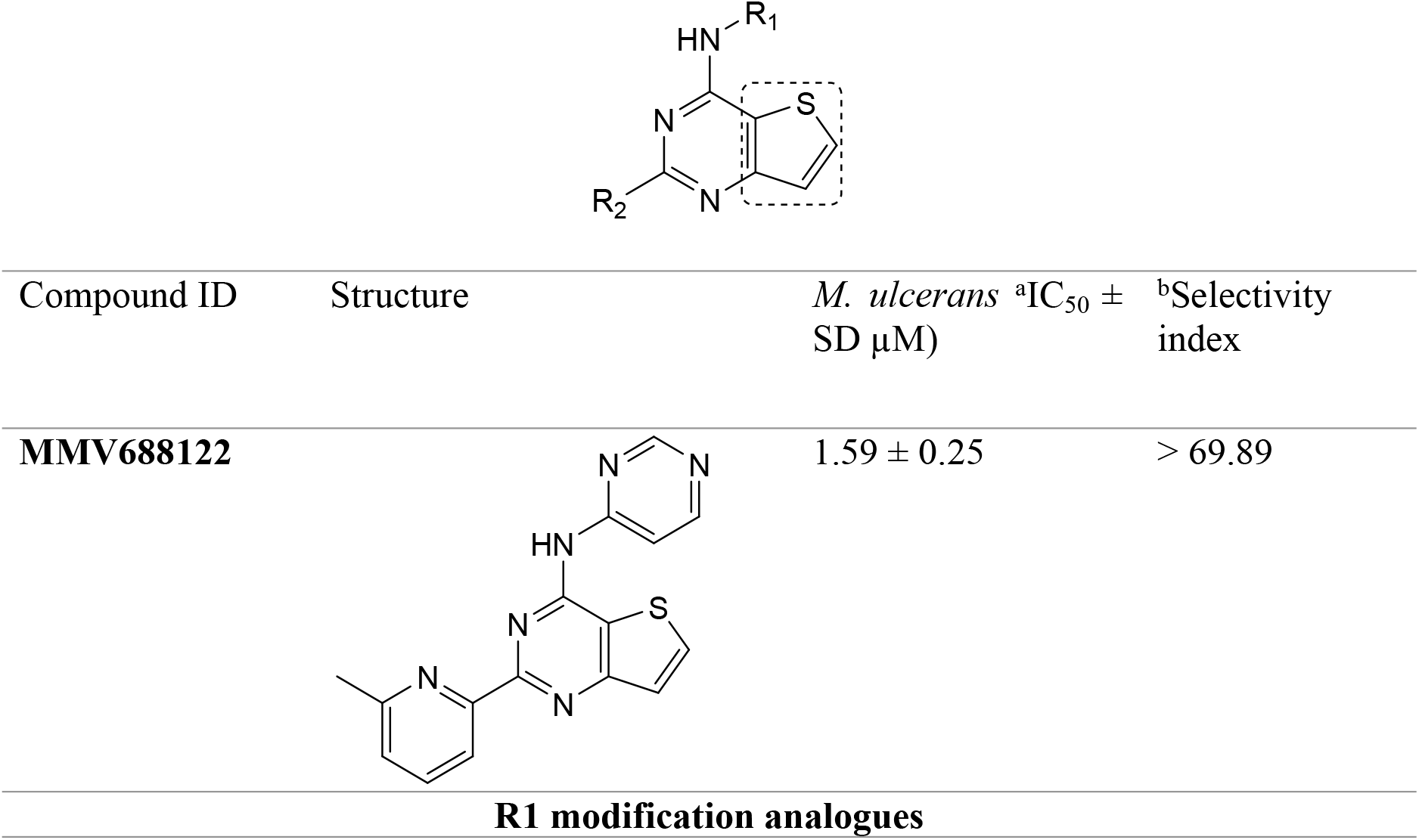
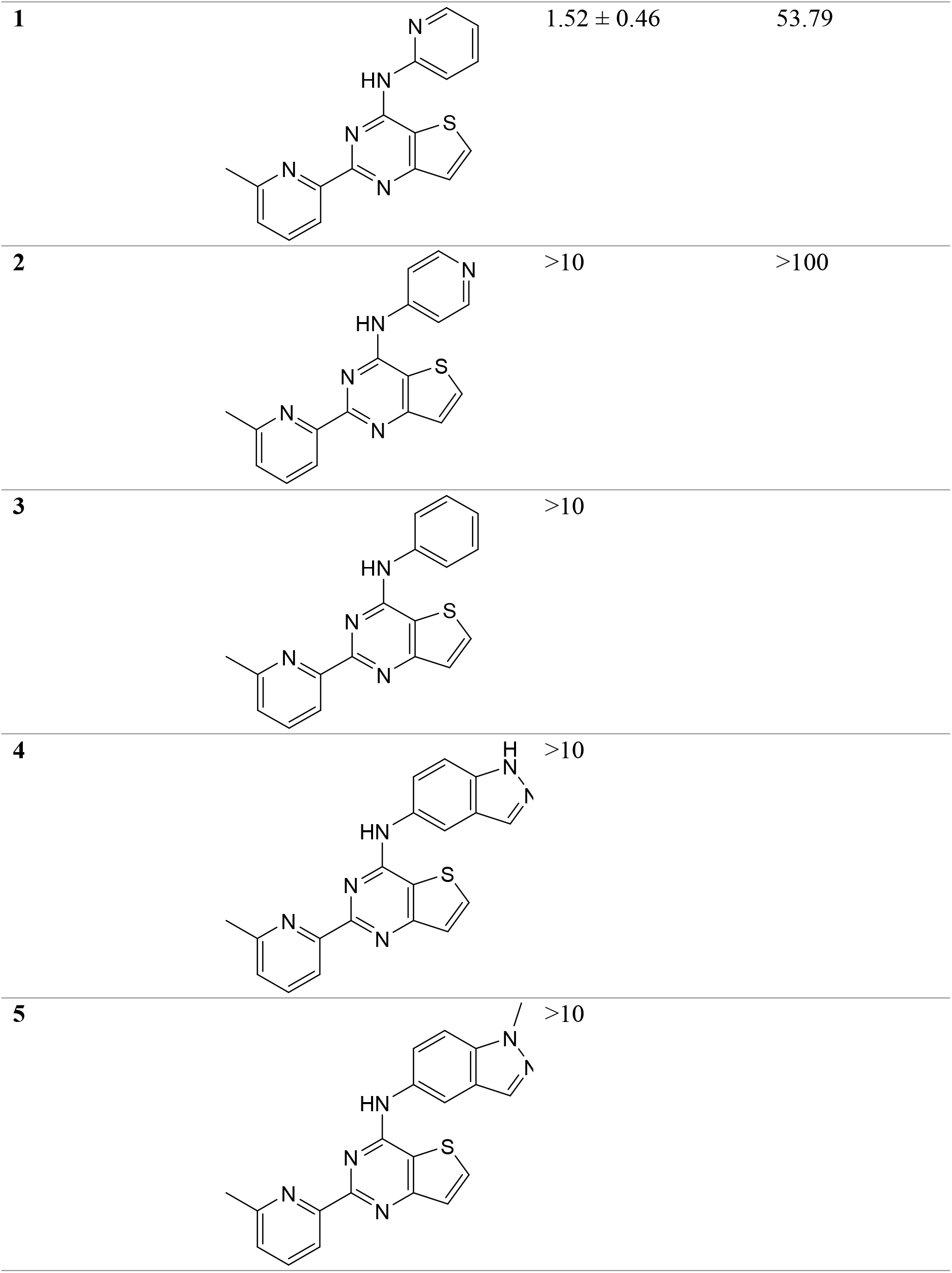
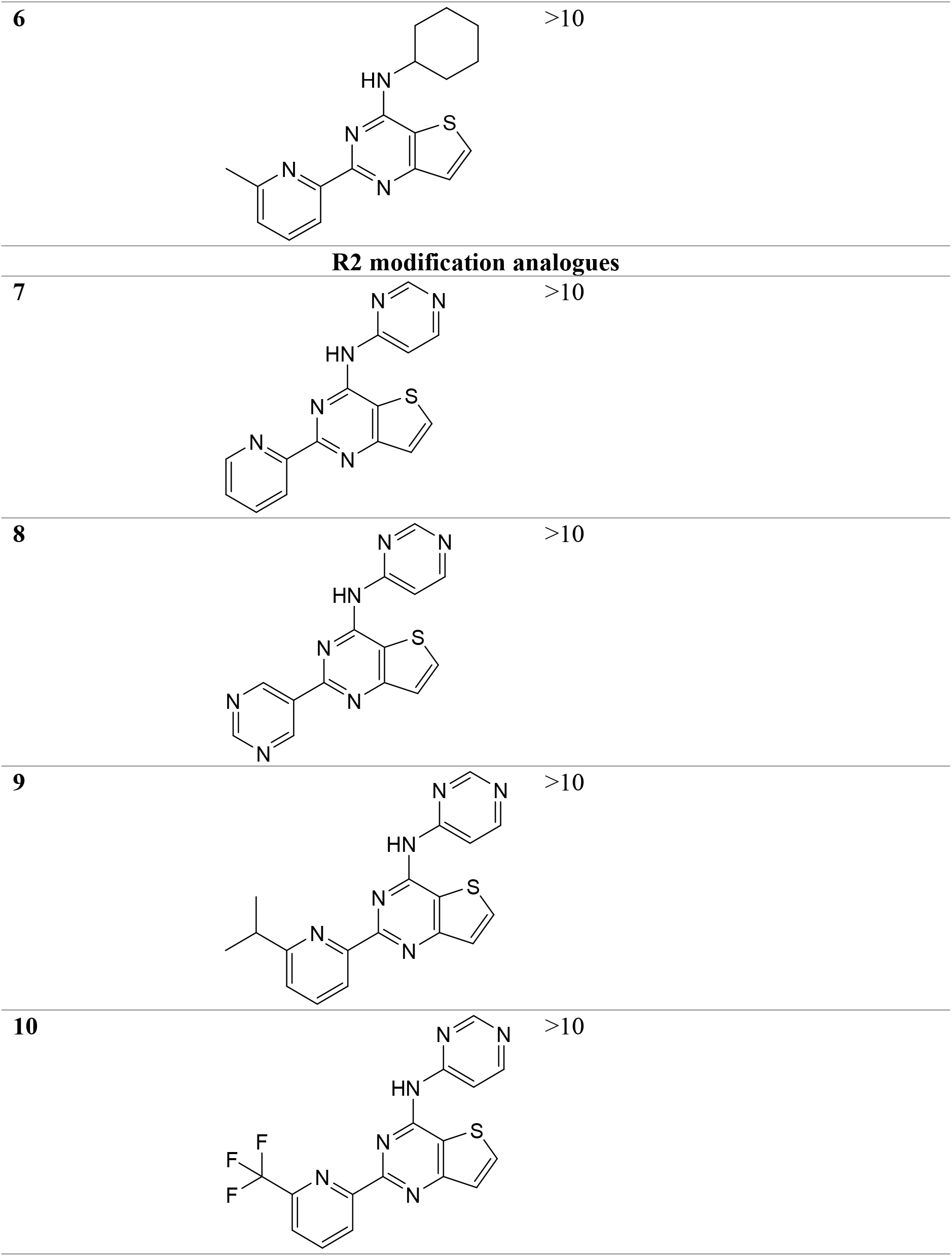
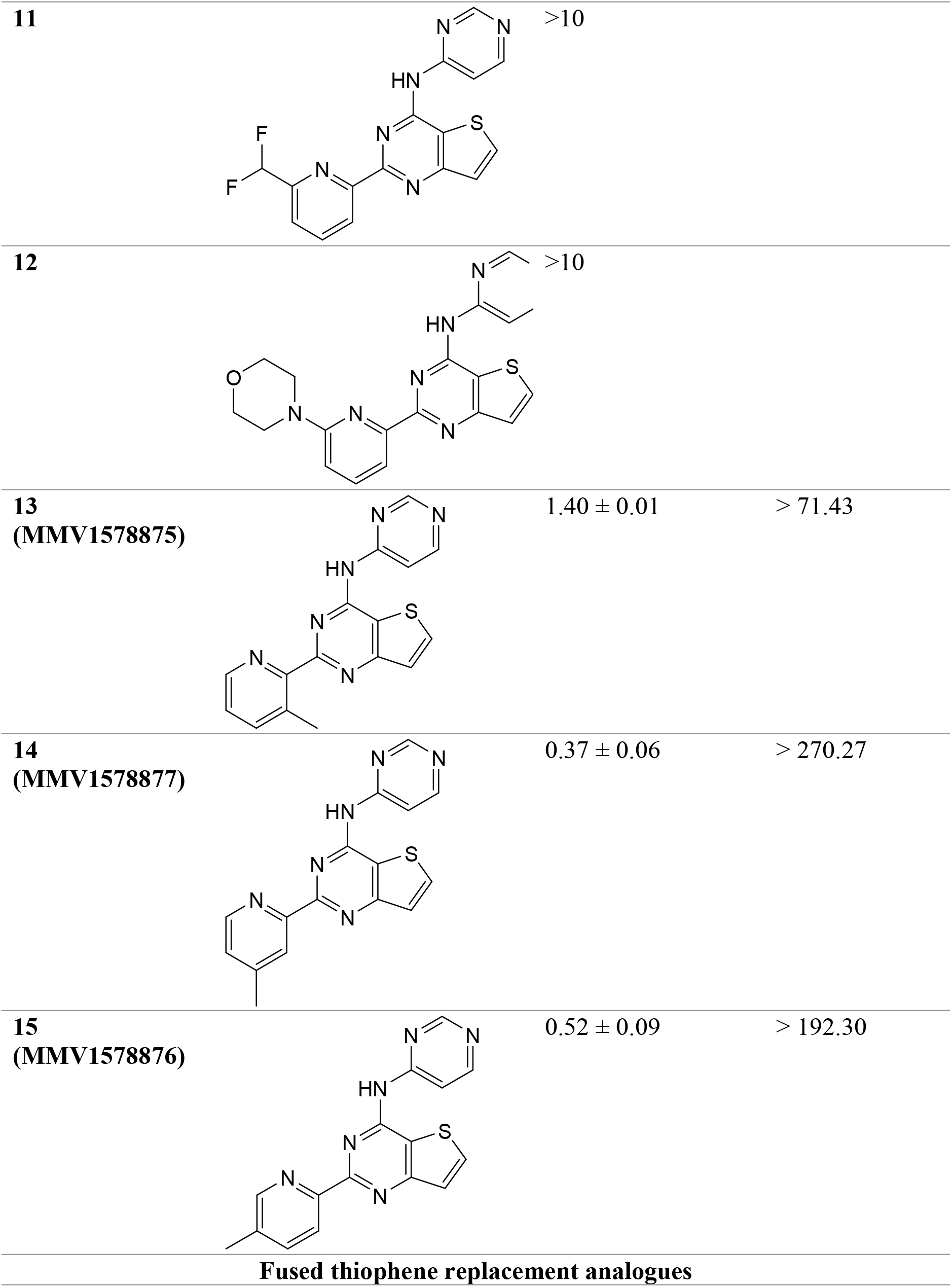
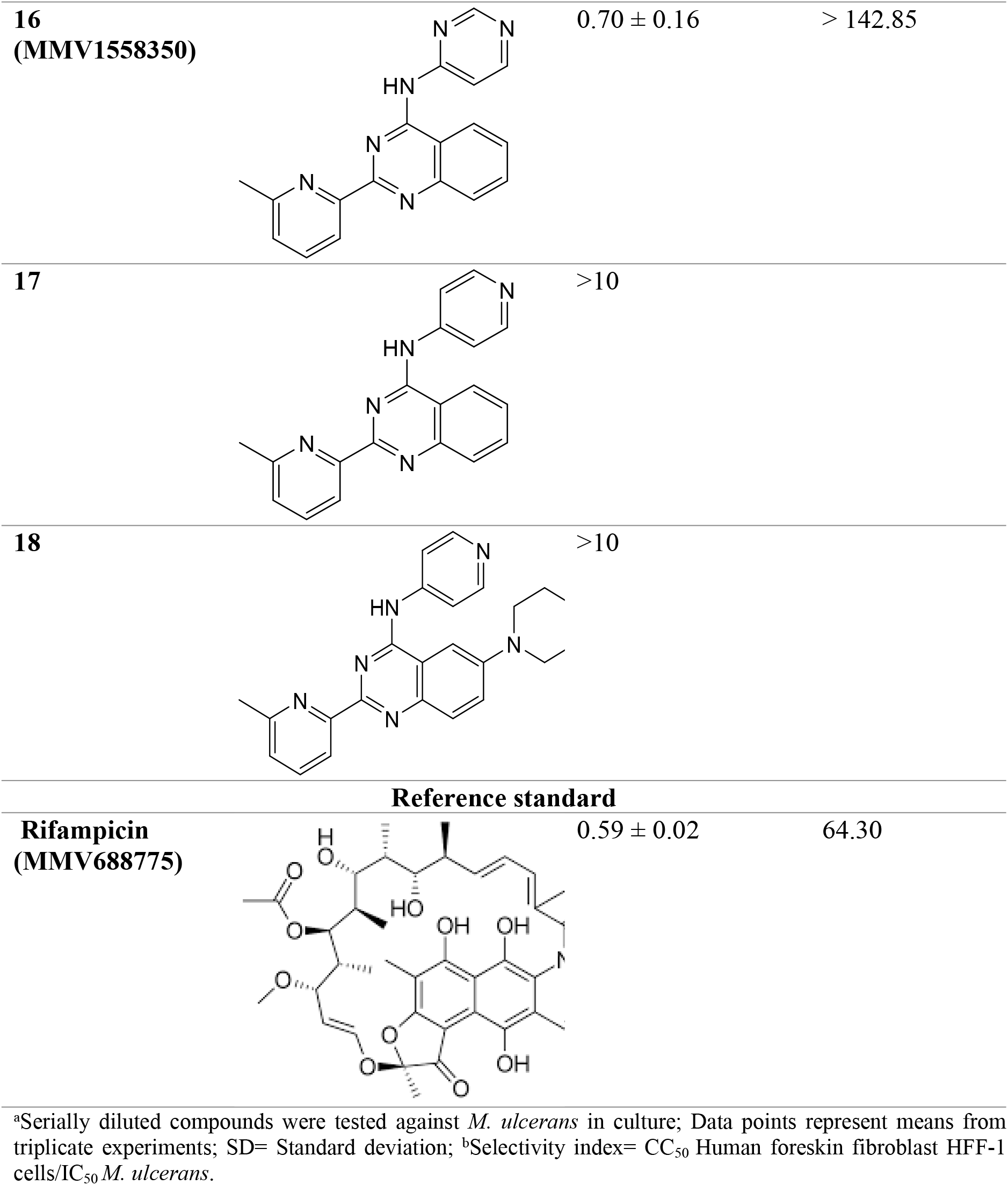
SAR studies on representative analogues of MMV688122.

The other 33 hits notoriously turned inactive and were considered as false positives. Compounds MMV688756 and MMV688327 belong to the oxazolidinone class [14]. In addition to their antituberculous activity, these compounds were found to inhibit the growth of nontuberculous *Mycobacteria*, including *M. abscessus* and *M. avium* [17]. Recently, sutezolid was shown to be more effective than linezolid, tedizolid, and delpazolid with respective minimum inhibitory concentrations (MIC) of 0.03 µg/mL, 1.13 µg/mL, 1.13 µg/mL, and 0.25 µg/mL when tested against clinical isolates of *M. ulcerans* isolated in Beijing, China [18]. However, although the anti-*M. ulcerans* activity of sutezolid is well established, the activity of its derivative radezolid has not yet been reported. To our knowledge, this is the first report on the antimycobacterial activity of this compound (IC_50_: 1.60 µM) against *M. ulcerans*. Furthermore, although the most promising novel chemotype MMV688122 (IC_50_: 1.59 µM) was previously shown to possess antituberculous activity [15], its anti-*M. ulcerans* activity is been reported here for the first time. Thus, MMV688122 (2-(6-methylpyridin-2-yl)-N-(pyrimidin-4-yl)thieno[3,2-d]pyrimidin-4-amine) represents a promising novel anti-*M. ulcerans* chemotype warranting further investigation.

### Structure-activity relationship (SAR) study of MMV688122 derivatives

The interesting anti-*M. ulcerans* activity of compound MMV688122 (IC_50_: 1.59 µM) prompted us to synthesize analogues to enable SAR studies. Structural analogues were synthesized and tested for anti-*M. ulcerans* activity. Results on 18 representatives are reported in Table 2. Among these compounds, five (MMV1578870, MMV1578875, MMV1578876, MMV1578877, and MMV1558350) were found to be more active (IC_50_ range: 0.37-1.52 μM) than the original hit compound MMV688122 (IC_50_: 1.59 μM). Compounds MMV1578876 (IC_50_ of 0.52 μM) and MMV1578877 (IC_50_ of 0.37 μM) exhibited comparable IC_50_ values when compared with the standard anti-*M. ulcerans* drug rifampicin (IC_50_: 0.59 μM) tested in the same conditions. When tested for cytotoxicity on human foreskin fibroblast HFF-1 cells, compound MMV1578877 (SI> 270.27) was ~1.6-fold more selective than rifampicin (SI: 169.5).

To study the SAR of MMV688122 and its derivatives, variations around the pyrimidine group (R1), the pyridine (R2), and the thiophene (core) were examined. First, replacement of the 2,4-pyrimidine group at the R1 part of the parent molecule by 2-pyridine led to compound 1 with similar potency (IC_50_ 1.52 µM). However, replacement with 4-pyridine, indazoles or cyclohexyl were not tolerated (compounds 2-6). Both removal of the methyl substituent (compound 7) and replacement by a 3,5-pyrimidine (compound 8) at the R2 part resulted in loss of activity (IC_50_s >10 μM). Given the apparent importance of the methyl group, alternative substituents were investigated: isopropyl, trifluoromethyl, difluoromethyl, and morpholine (compounds 9-12). As none of these changes appeared to be tolerated, the position of the methyl group of the pyridine skeleton (R2) was examined if important for the activity against *M. ulcerans*. Changes in the methyl position (3, 4 and 5 of the pyridin-2-yl pharmacophoric moiety) for compounds 13 (MMV1578875), 14 (MMV1578877) and 15 (MMV1578876), respectively affected the antimycobacterial activity (IC_50_ values: 1.40, 0.37, 0.52 μM, respectively) compared to the parent counterpart (MMV688122; 6-methylpyridin-2-yl skeleton; IC_50_: 1.59 μM). These findings demonstrate the strong implication of methylpyridin-2-yl pharmacophoric moiety in inhibiting the growth of *M. ulcerans*.

To explore further the SAR, the core moiety was investigated, especially the fused thiophene skeleton due to its potential risk related to the formation of reactive metabolites. Noteworthily, the replacement of the fused thiophene group in MMV688122 (IC_50_ of 1.59 μM) by a fused benzene ring improved the activity by 2-fold (IC_50_ of 0.70 μM for compound 16, MMV1558350), suggesting that quinazoline could be a more promising core moiety than the thienopyrimidine for antimycobacterial activity. Lastly, the lack of nitrogen at position 5 within the pyrimidine ring of compounds 17 and 18 (IC_50_s >10 μM) confirmed the loss of the antimycobacterial activity previously observed with the fused thiophene core (compound 2).

The *in vitro* DMPK (drug metabolism and pharmacokinetics) and physicochemical properties of this chemical series were then explored. Representative derivatives were tested for oxidative metabolism in human and mouse liver microsomes, and mouse hepatocytes. Distribution coefficient (LogD) and aqueous solubility were also measured (Table 3). All tested compounds showed poor metabolic stability in both human and mouse species. Interestingly, compound 13 (MMV1578875) exhibited the highest microsomal stability in human microsomes (36 µL/min/mg protein). On the other hand, all tested thienopyrimidine derivatives demonstrated good aqueous solubility. Lower solubility was observed for the quinazoline representative (compound 16, MMV1558350) but remained in an acceptable range. This can be explained by the higher LogD associated with this core moiety.

**Table 3:**
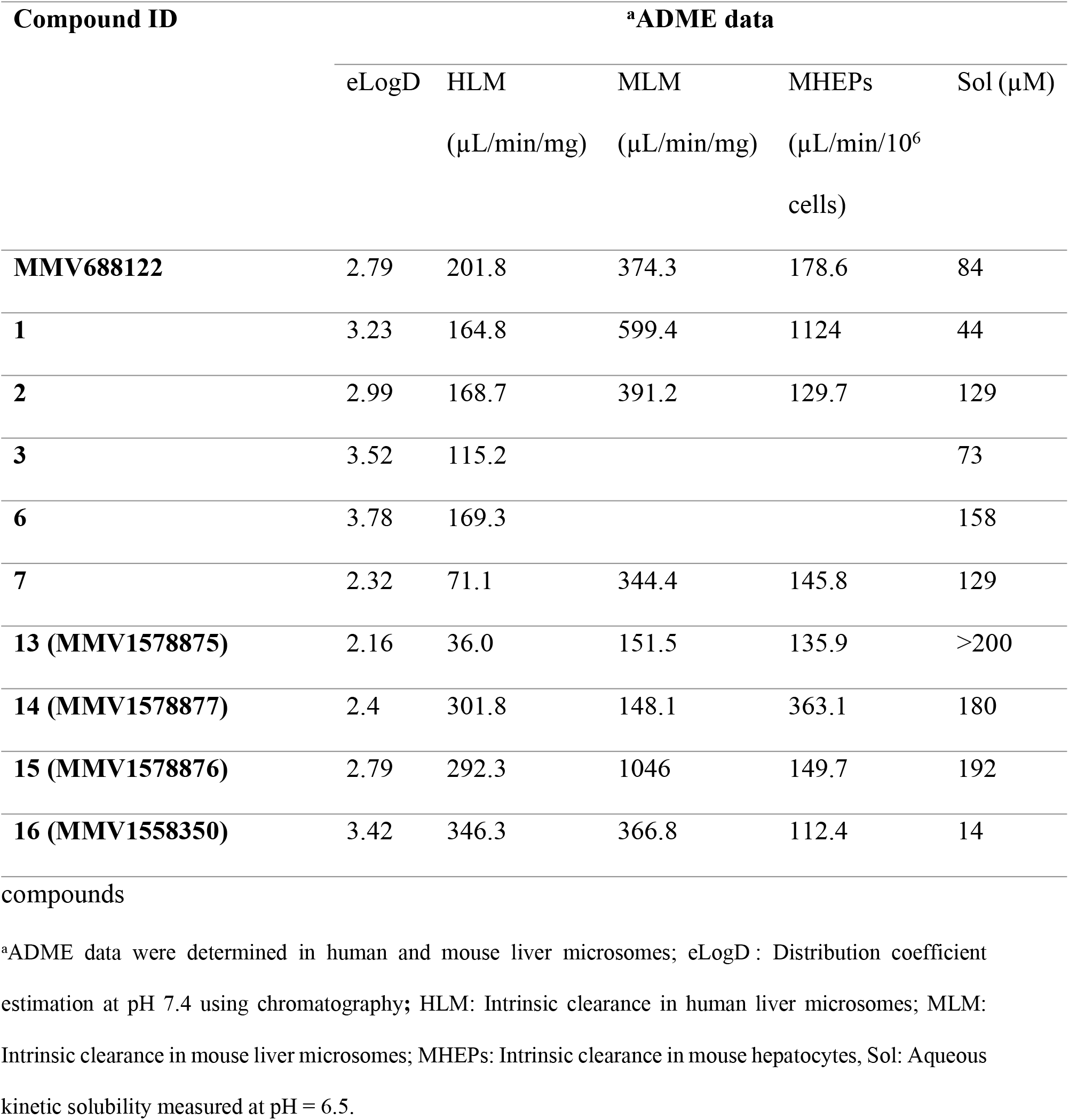
*In vitro* DMPK and physicochemical properties of selected anti-*M. ulcerans* hit.

| Compound ID | <sup>a</sup>ADME data |  |  |  |  |
| --- | --- | --- | --- | --- | --- |
| | eLogD | HLM<br>($\mu\text{L}/\text{min}/\text{mg}$) | MLM<br>($\mu\text{L}/\text{min}/\text{mg}$) | MHEPs<br>($\mu\text{L}/\text{min}/10^6$<br>cells) | Sol ($\mu\text{M}$) |
| **MMV688122** | 2.79 | 201.8 | 374.3 | 178.6 | 84 |
| **1** | 3.23 | 164.8 | 599.4 | 1124 | 44 |
| **2** | 2.99 | 168.7 | 391.2 | 129.7 | 129 |
| **3** | 3.52 | 115.2 |  |  | 73 |
| **6** | 3.78 | 169.3 |  |  | 158 |
| **7** | 2.32 | 71.1 | 344.4 | 145.8 | 129 |
| **13 (MMV1578875)** | 2.16 | 36.0 | 151.5 | 135.9 | >200 |
| **14 (MMV1578877)** | 2.4 | 301.8 | 148.1 | 363.1 | 180 |
| **15 (MMV1578876)** | 2.79 | 292.3 | 1046 | 149.7 | 192 |
| **16 (MMV1558350)** | 3.42 | 346.3 | 366.8 | 112.4 | 14 |

compounds
<sup>a</sup>ADME data were determined in human and mouse liver microsomes; eLogD : Distribution coefficient estimation at pH 7.4 using chromatography; HLM: Intrinsic clearance in human liver microsomes; MLM: Intrinsic clearance in mouse liver microsomes; MHEPs: Intrinsic clearance in mouse hepatocytes, Sol: Aqueous kinetic solubility measured at pH = 6.5.

### Metabolite identification studies on MMV1578877

The most potent compound, MMV1578877 (compound 14), was selected for metabolite identification studies. The *in vitro* incubation of MMV1578877 was conducted at 10 µM substrate and 1 mg/mL microsomal protein to facilitate metabolite detection. Degradation of parent compound was only detected in the presence of NADPH suggesting that the loss was mediated by CYP enzymes. Three putative metabolites were detected; two with masses consistent with mono-oxygenation (M+16 (I) and (II), m/z 337) and one di-oxygenated product (M+32, m/z 353). Based on MS/MS analysis, the site of oxygenation is likely at the methylpyridine moiety for all three metabolites. Relative peak areas suggested that the most abundant metabolite in human liver microsomes was a mono-oxygenation product whereas in mouse microsomes, the most abundant metabolite appeared to be a di-oxygenation product. Authentic metabolite standards would be required for structural confirmation and quantification.

## Conclusion

Screening of the Pathogen Box allowed identification of four compounds (MMV688122, MMV688327, MMV688756 and MMV688775) with low IC_50_ values (0.59 to 3.34 µM) against *Mycobacterium ulcerans* and high selectivity toward HFF cells (CC_50_ > 100 µM). Compounds MMV688327, MMV688756 and MMV688775 correspond to radezolid, sutezolid and rifampicin (standard antimycobacterial drug), respectively. To our knowledge, this is the first report on the antimycobacterial activity of radezolid (IC_50_ 1.60 µM) against *Mycobacterium ulcerans* and for the 2-(pyridin-2-yl)-N-(pyrimidin-4-yl)thieno[3,2-d]pyrimidin-4-amine MMV688122 (IC_50_ 0.59 µM, SI > 69.89). SAR studies were performed on the latter chemotype thanks to the synthesis of structural analogues. Among these, MMV1578877 was found to be the most potent (IC_50_ 0.37 µM, SI >270.27). The compounds from this thienopyrimidine class present good aqueous solubility but suffer from poor metabolic stability. Therefore, further optimization through medicinal chemistry is required to improve pharmacokinetic properties to enable *in vivo* studies. The improved metabolic stability observed for MMV1578875 in human liver microsomes and the possibility to move towards quinazoline as core moiety (MMV1558350) whilst maintaining attractive potency constitute promising findings to develop this novel chemotype as anti-*M. ulcerans* agent.

## Acknowledgements

The authors are very grateful to Medicines for Malaria Venture (MMV), which provided the open access Pathogen Box and structural analogues. The metabolite identification studies were conducted at the Centre for Drug Candidate Optmisation, Monash University under the direction of Prof. Susan Charman.

**S1: Experimental procedures for the synthesis of MMV688122 and compounds 1-18**

Synthesis and characterization of MMV688122 and compounds 1-6

Synthesis and characterization of compounds 7-10 and compounds 12-15

Synthesis and characterization of compounds 16 and 17:

Synthesis and characterization of compound 18

**S2: 1HNMR Spectra and Chromatograms of Compounds 1-18**

